# A novel chimeric RNA originating from BmCPV S4 and *Bombyx mori* HDAC11 transcripts regulates virus proliferation

**DOI:** 10.1101/2023.02.07.527451

**Authors:** Jun Pan, Shulin Wei, Qunnan Qiu, Xinyu Tong, Zeen Shen, Min Zhu, Xiaolong Hu, Chengliang Gong

## Abstract

Polymerases encoded by segmented negative-strand RNA viruses cleave 5’-m7G-capped host transcripts to prime viral mRNA synthesis (‘‘cap-snatching’’) to generate chimeric RNA, and trans-splicing occurs between viral and cellular transcripts. *Bombyx mori* cytoplasmic polyhedrosis virus (BmCPV), an RNA virus belonging to Reoviridae, is a major pathogen of silkworm (*B. mori*). The genome of BmCPV consists of 10 segmented double-stranded RNAs (S1-S10) from which viral RNAs encoding a protein are transcribed. In this study, chimeric silkworm-BmCPV RNAs, in which the sequence derived from the silkworm transcript could fuse with both the 5’ end and the 3’ end of viral RNA, were identified in the midgut of BmCPV-infected silkworms by RNA_seq and further confirmed by PCR and Sanger sequencing. A novel chimeric RNA, HDAC11-S4 RNA 4, derived from silkworm histone deacetylase 11 (HDAC11) and the BmCPV S4 transcript encoding viral structural protein 4 (VP4), was selected for validation by *in situ* hybridization and Northern blotting. Interestingly, our results indicated that HDAC11-S4 RNA 4 was generated in a BmCPV RNA-dependent RNA polymerase (RdRp)-independent manner and could be translated into a truncated BmCPV VP4 with a silkworm HDAC11-derived N-terminal extension. Moreover, it was confirmed that HDAC11-S4 RNA 4 inhibited BmCPV proliferation, decreased the level of H3K9me3 and increased the level of H3K9ac. These results indicated that during infection with BmCPV, a novel mechanism, different from that described in previous reports, allows the genesis of chimeric silkworm-BmCPV RNAs with biological functions.

**Graphical abstract:** 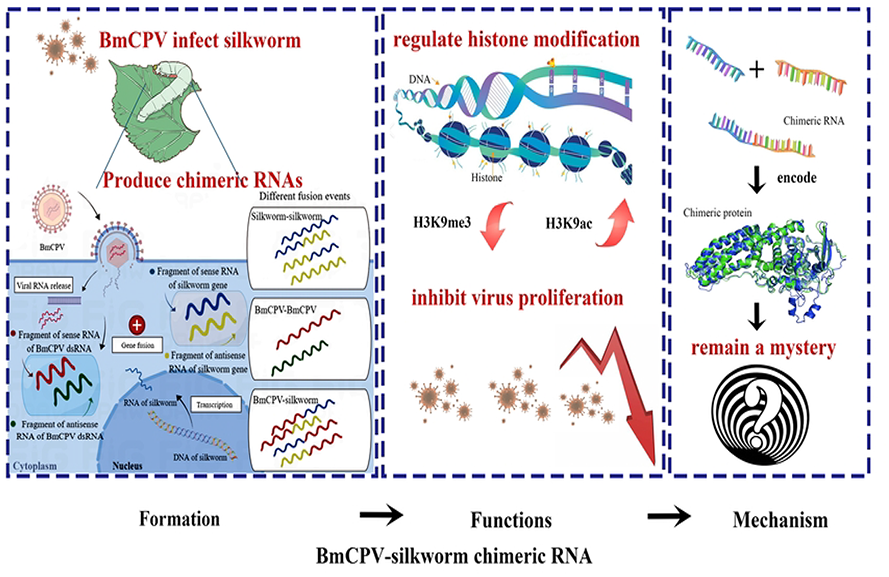

---

Most eukaryotic mRNAs have a 5’ methyl-7-guanosine (m7G) cap structure that recruits ribosomes to initiate translation in the cytoplasm(Decroly et al., 2011; Leppek et al., 2018). Segmented negative-strand RNA viruses (sNSVs), lacking their own capping machinery, cannot synthesize the 5’ terminal m^7^G cap structure for their mRNAs, but the capped host transcripts can be cleaved, resulting in the generation of a capped primer. Viral mRNA synthesis is initiated by using the capped primer. This process has been referred to as “cap-snatching”(Rialdi et al., 2017). The 5’-end of chimeric host and virus mRNAs generated via the cap-snatching mechanism have highly diversified sequences derived from the 5’-ends of host mRNAs(Sikora et al, 2017; Clohisey et al, 2020). Unlike canonical cap-snatching, influenza A virus (IAV) can utilize noncanonical cap-snatching to diversify its mRNAs/ncRNAs(Li et al., 2020). Once the host translational machinery is generated, it can be recruited to host-viral mRNAs for translation, which results in global transcriptional suppression in the host and evasion of antiviral surveillance by RIG-I-like receptors(Decroly et al., 2011; Ho et al., 2020; Russell, 2020).

It has been reported that start codons within cap-snatched host transcripts can generate chimeric host-viral mRNAs with coding potential. sNSVs can obtain functional upstream start codons (uAUGs), via a process termed “start-snatching”(Ho et al., 2020).

Translation is initiated from host-derived uAUGs in chimeric host-viral transcripts, which results in the generation of two novel chimeric types of proteins depending on the open reading frame (ORF) of the uAUG relative to that of the canonical viral protein: canonical viral proteins with N-terminal extensions derived from host and viral untranslated regions (UTRs) and novel proteins that are translated from ORFs differing from canonical viral ORFs(Ho et al., 2020). Previous studies suggested that novel viral proteins generated by cap-snatching impact viral replication and serve as additional targets for host surveillance (Russell, 2020; Ho et al., 2020).

To date, the cap-snatching mechanism has been found in sNSVs belonging to the Peribunyaviridae, Phenuiviridae, Tospoviridae, Arenaviridae, Nairoviridae, Hantaviridae, and Orthomyxoviridae families(Ho et al., 2020; Olschewski et al., 2020).

Two distinct RNA transcripts can be joined by trans-splicing, resulting in the formation of chimeric RNA. A previous study showed naturally occurring heterologous trans-splicing of adenovirus (ADV) (linear double-stranded DNA virus) RNA with host cellular transcripts during infection (Kikumori et al., 2002). Moreover, it has been found that heterologous trans-splicing also occurs between human immunodeficiency virus (HIV, single-stranded RNA virus)-nef RNA and cellular transcripts (Caudevilla et al., 2001), and a 100 kDa super T antigen harboring a duplication of the retinoblastoma (pRb)-binding domain can be generated by homologous RNA trans-splicing of viral early transcripts during infection with simian virus 40 (SV40, cyclic double-stranded DNA virus) (Eul et al., 2013). Chimeric RNAs can also be generated by heterologous SV40 transcript trans-splicing, but their function is unknown (Poddar et al., 2014). A recent study showed that novel proteins associated with the viral life cycle were produced by trans-splicing of late transcripts of human polyoma JC virus (JCV, cyclic double-stranded DNA virus) (Saribas et al., 2018). In addition to trans-splicing, it has been reported that a functional mRNA encoding a fusion of the viral E3 ubiquitin ligase ICP0 and viral membrane glycoprotein L can be produced by low level readthrough transcription during herpes simplex virus type 1 (HSV-1) infection (Depledge et al. 2019) and that chimeric cellular-HIV mRNAs can be generated by aberrant splicing (Lee et al., 2022).

Cytoplasmic polyhedrosis viruses (CPVs) with segmented double-stranded RNAs (dsRNAs) that are packaged into a single-layered icosahedral viral capsid (Zhang et al., 2015) belong to the *Cypovirus* genus of the Reoviridae family. CPVs can infect insects belonging to Lepidoptera, Hymenoptera and Coleoptera and play a very important role in the control of pest populations in agriculture and forestry. *Bombyx mori* CPV (BmCPV), a model CPV species, is a pathogen of silkworm, *B. mori*, and specifically infects epithelial cells of the midgut, resulting in a decrease in cocoon production (Arella et al., 1988). The BmCPV genome consists of 10 dsRNA segments(Patton and Spencer, 2000). Previous studies have indicated that the viral structural proteins VP1, VP2, VP3, VP4, VP6 and VP7 are encoded by the S1, S2, S3, S4, S6 and S7 segments, respectively, and that the nonstructural proteins NSP5, NSP8, NSP9 and polyhedrin are encoded by the S5, S8, S9 and S10 segments (Cao et al., 2012). The transcript of each segment possesses a 5’-m7G cap structure (Smith & Furuichi, 1982), which is considered a monocistron encoding a protein. In recent decades, it was widely believed that viral transcripts were not spliced during the formation of mature viral mRNAs. However, recent studies have indicated that BmCPV RNAs can be cut by Dicer-2 and an uncharacterized endo-RNase to produce thousands of small viral RNAs (vsRNA) as a strategy for opposing virus infection (Zografidis et al., 2015), and transcripts of BmCPV can also be processed to viral microRNAs (vmiRNAs) to promote virus infection (Zhao et al., 2022; Li et al., 2021; Wang et al., 2021; Guo et al., 2020). Our previous studies indicated that BmCPV RNAs can be processed to form viral circular RNAs (vcircRNAs) with an unknown mechanism. Both circRNA-vSP27 and cirRNA 000048, derived from BmCPV, can attenuate viral replication because they encode the small peptides vSP27 and vsp21 (Zhang et al., 2022a; 2022b), respectively. Moreover, a BmCPV S7 RNA-derived circular DNA (vcircDNA) referred to as vcDNA-S7 was detected in infected cells, and vcDNA-S7 was transcribed into RNA, which was further processed into antiviral vsRNAs (Zhu et al., 2022). These results indicated that functional molecules, including vsRNAs, vmiRNAs, vcircRNAs and vcircDNA, can be formed from BmCPV RNAs. However, whether chimeric silkworm-BmCPV RNAs can be generated during BmCPV infection remains a mystery.

In this study, we identified chimeric silkworm-BmCPV RNAs and identified a novel chimeric RNA, HDAC11-S4 RNA 4, derived from silkworm histone deacetylase 11 (HDAC11), and BmCPV S4 RNA transcripts were shown to be generated in a BmCPV RNA-dependent RNA polymerase (RdRp)-independent manner. Moreover, we confirmed that a truncated BmCPV VP4 with a silkworm HDAC11-derived N-terminal extension, translated by HDAC11-S4 RNA 4, could inhibit BmCPV proliferation, decrease the level of H3K9me3 and increase the level of H3K9ac. These results indicated that a novel mechanism, different from that described in previous reports, allows the genesis of chimeric silkworm-BmCPV RNAs with biological functions.

## 1 Results

### 1.1 Chimeric silkworm-BmCPV RNAs can be formed during BmCPV infection

It has been found that chimeric host‒virus RNAs can be formed by cap-snatching during sNSV and HIV infection (Ho et al., 2020; Caudevilla et al., 2001) or by trans-splicing during dsDNA virus infection (Eul et al., 2013; Saribas et al., 2018); however, chimeric host-viral RNAs have not been found during dsRNA virus infection thus far. To explore whether chimeric silkworm-BmCPV RNA can be formed, the BmCPV-infected midgut was subjected to RNA_Seq. The raw sequencing data have been uploaded to the NCBI database (accession numbers SRR22891215, SRR22891214, SRR22891213). After removing low-quality reads, the clean reads were assembled with StringTie software (1.2.0) (Pertea et al., 2015) using the *B. mori* genome and the BmCPV genome as references. STAR-Fusion was used to analyze fusion transcripts (Haas et al., 2019). A total of 516 redundant chimeric silkworm RNAs were identified, and changes in the number and abundance of chimeric transcripts were found following virus infection. Interestingly, 372 redundant chimeric viral RNAs and 34 redundant chimeric silkworm-viral RNAs (Table S1) were found in the BmCPV-infected midgut (Figure 1). Most of the chimeric silkworm-silkworm RNAs were derived from two different transcripts, but some chimeric silkworm-silkworm RNAs were formed by the fusion of 3 fragments from different transcripts (Figure 1). Among the chimeric virus‒virus RNAs, all detected chimeric RNAs originated from fusions between different BmCPV RNA fragments, which were derived from either the fusion of sense strands of viral RNA fragments or the fusion of antisense strands of viral RNA fragments (Figure 1). Among the chimeric silkworm-BmCPV RNAs, chimeric RNAs derived from the fusion of sense/antisense RNA fragments of silkworm genes and sense/antisense RNA fragments of BmCPV genomic dsRNAs were identified in the BmCPV-infected midgut. Silkworm RNA sequences can be fused with the 5’ or 3’ terminus of a viral RNA fragment (Figure 1). Among the chimeric RNAs, the most frequently detected RNA fragment derived from silkworm transcripts was large subunit ribosomal RNA (XR_005246581.1), followed by the mRNA clone fcaL43P13 (AK384927.1), U6 atac minor spliceosomal RNA (XR_005245887.1), Bm_160 RNA (GU247410.1) and histone deacetylase (HDAC) 11 (XM_004925365.4). The flanking sequences of the junction sites of the selected chimeric RNAs were identified by PCR and Sanger sequencing, and the results were consistent with those of high-throughput sequencing, suggesting that the results obtained from high-throughput sequencing were credible (Figure 2).

**Figure 1.**
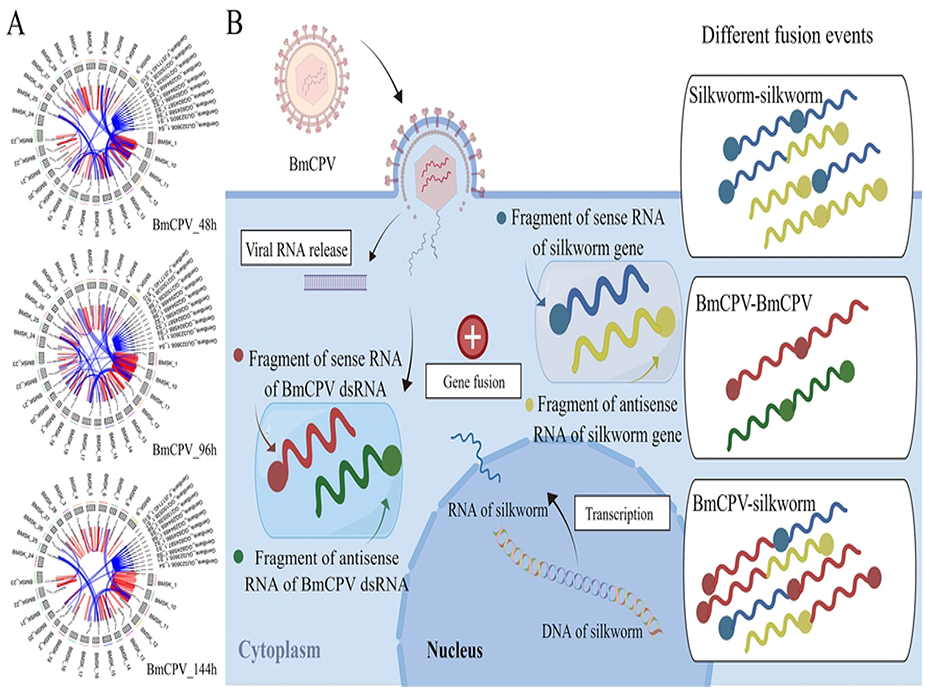
Chimeric RNAs generated in the silkworm midgut infected with BmCPV. **A**, Circos diagram of chimeric RNAs in the midgut at 48 h (BmCPV_48), 96 h (BmCPV_96) and 144 h (BmCPV_144) postinfection with BmCPV; **B**, Fusion types of chimeric RNAs detected in the midgut infected with BmCPV.

**Figure 2.**
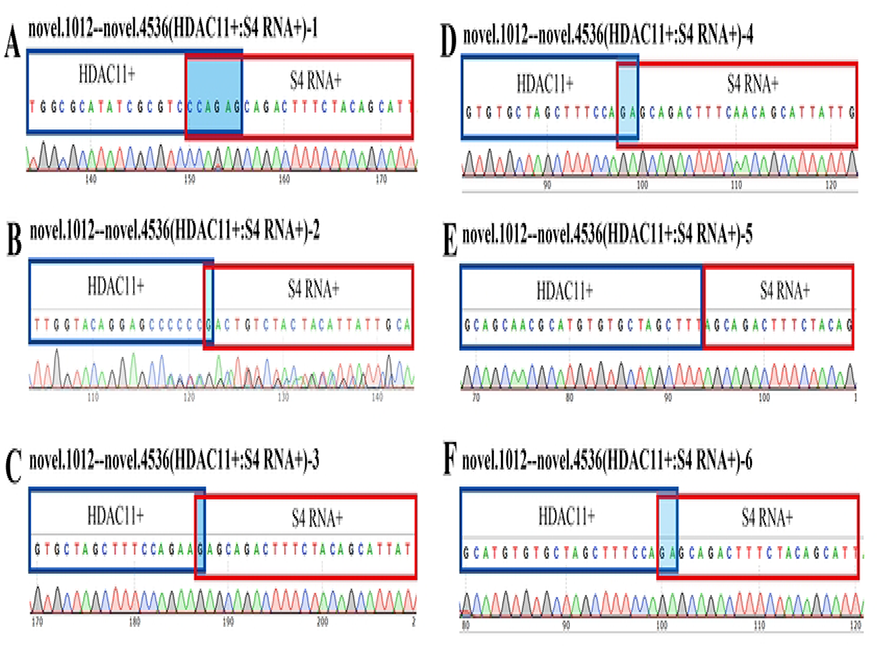
Identification of chimeric RNAs by PCR and Sanger sequencing. **A and B**, fragments from the antisense RNAs of BmCPV S3 (A) and S4 (B) dsRNAs fused with a fragment of the larger subunit of ribosomal RNA (LSrRNA) from silkworm, the sequence indicated with a blue background is a common sequence shared by the S3/S4 and LSrRNA antisense sequences. **C**, Fragment of antisense RNA of BmCPV S7 dsRNA fused with a fragment of the silkworm mRNA clone: fcaL43P13. **D**, Fragment of silkworm U6 ncRNA fused with a fragment of sense RNA BmCPV S7 dsRNA. **E and F**, a fragment of silkworm U6 ncRNA/Bm_160 RNA fused with a fragment of the sense RNA of BmCPV S9 dsRNA. **G**, Fragment of antisense RNA of BmCPV S7 dsRNAs fused with a fragment of the antisense RNA of BmCPV S5 dsRNAs. **H**, A fragment of the sense RNA of BmCPV S9 dsRNAs fused with a fragment of the sense RNA of BmCPV S7 dsRNAs. The sequences indicated with a blue background are sequences shared by two parental RNAs.

### 1.2 Chimeric silkworm-BmCPV RNAs are formed in a different way than previously reported

The trans-splicing of viral transcripts with host or viral transcripts can occur in the nucleus of infected cells through a splicing mechanism (Caudevilla et al., 2001; Eul et al., 2013; Poddar et al., 2014; Saribas et al., 2018; Lee et al., 2022). In this study, we defined the parental RNAs of the sequences located upstream/downstream of the fusion site of chimeric silkworm-BmCPV RNAs as left/right RNAs. To understand the formation mechanisms of chimeric RNAs, we analyzed the flanking sequences of the breakpoints of the parental RNAs, and the conserved sequences required for the splicing mechanism were not found (Figure S1). The transcription of BmCPV RNAs occurred in the cytoplasm, suggesting that chimeric silkworm-BmCPV RNAs were formed in a manner different from trans-splicing.

Cap-snatching is a mechanism applied by sNSVs to initiate genome transcription (Xu et al., 2022). The 5’-ends of generated chimeric host and virus mRNAs have highly diversified sequences derived from the 5’-ends of host mRNAs (Sikora et al, 2017; Clohisey et al, 2020). In this study, chimeric silkworm-BmCPV RNAs were found to be formed by the fusion of sense/antisense RNA fragments of silkworm genes and sense/antisense RNA fragments of BmCPV genomic RNAs. A silkworm RNA sequence can be fused with the 5’ or 3’ terminus of a viral RNA fragment, implying that chimeric silkworm-BmCPV RNAs are formed in a manner different from cap-snatching (Figure 1).

Moreover, the fusion events between silkworm and BmCPV RNAs can be divided into two types. In one of these categories, the 5’-flanking sequence of the left parental RNA breakpoint and the 3’-flanking sequence of the right parental RNA breakpoint share a common sequence of 1-10 nt, and only one common sequence is retained in the formed chimeric RNA. In the other, two RNA fragments derived from different RNAs are directly joined after the RNAs are broken (Figure 2).

### 1.3 Isoforms of chimeric silkworm HDAC11-S4 RNA truly exist in the midgut infected with BmCPV

In this study, chimeric silkworm HDAC11-S4 RNA was identified in the BmCPV-infected midgut by high-throughput sequencing. To confirm this result, RT‒ PCR was carried out with two pairs of primers (HDAC11+:S4 RNA+)20-123 and (HDAC11+:S4 RNA+)20-456 (Table S2), which were designed according to the flanking sequences of the junction site of chimeric silkworm HDAC 11-S4 RNA. The sequencing results for the PCR products showed that 6 isoforms (HDAC11-S4 RNA 1-6) of HDAC 11-S4 RNA were identifiable, suggesting that the number of chimeric silkworm-BmCPV RNAs obtained by RNA_Seq was underestimated. With the exception of HDAC11-S4 RNA 2, the right flanking sequences of the junction sites of the remaining chimeric RNAs were from the 884-3262 nt region of the sense chain of S4 dsRNA (GU323606.1) (Figure 3).

**Figure 3.**
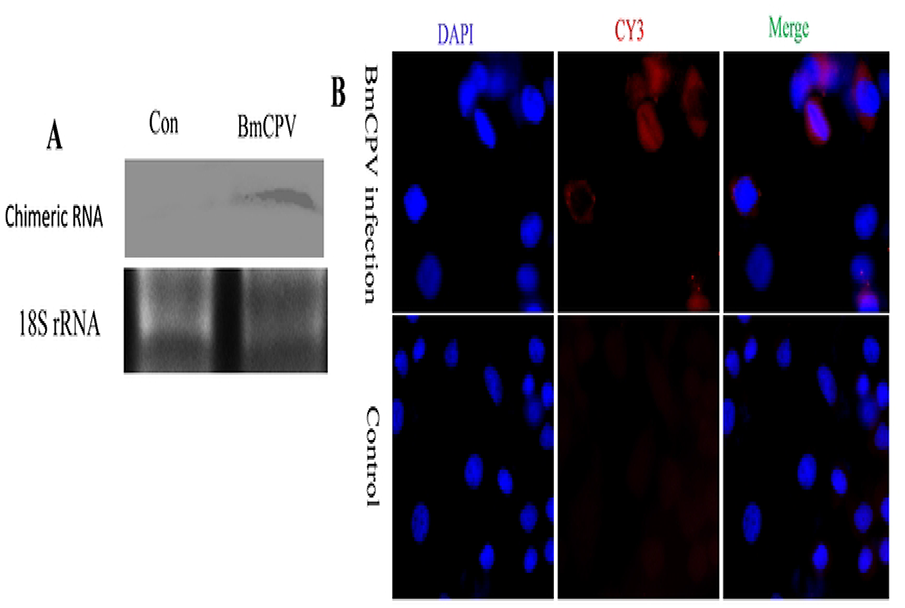
Isoforms of chimeric silkworm HDAC11-S4 RNA. A, B, C, D, E and F represent the junction sites of isoforms of chimeric silkworm HDAC11-S4 RNA.

To eliminate misjudgment caused by sequencing artifacts, reverse transcriptional noise, template-switching and scrambled junctions of RNAs, HDAC11-S4 RNA 4 was selected for further validation. PCR was conducted with the adapterHCPV-20-4 primer pair, in which one primer targeted the junction site of HDAC11-S4 RNA 4, and the other targeted the downstream sequence of the junction site of HDAC11-S4 RNA 4. The sequence of the PCR product was consistent with the expected result. The total RNAs extracted from the midgut infected with BmCPV were identified by Northern blotting with a biotin-labeled probe targeting the junction site of HDAC11-S4 RNA 4, and the results showed that a specific signal band was found in the RNAs extracted from the midgut infected with BmCPV (Figure 4A). Moreover, BmCPV-infected BmN cells were detected by *in situ* hybridization with the probe mentioned above. As expected, HDAC11-S4 RNA 4 was identified in the cytoplasm of infected cells (Figure 4B).

**Figure 4.**
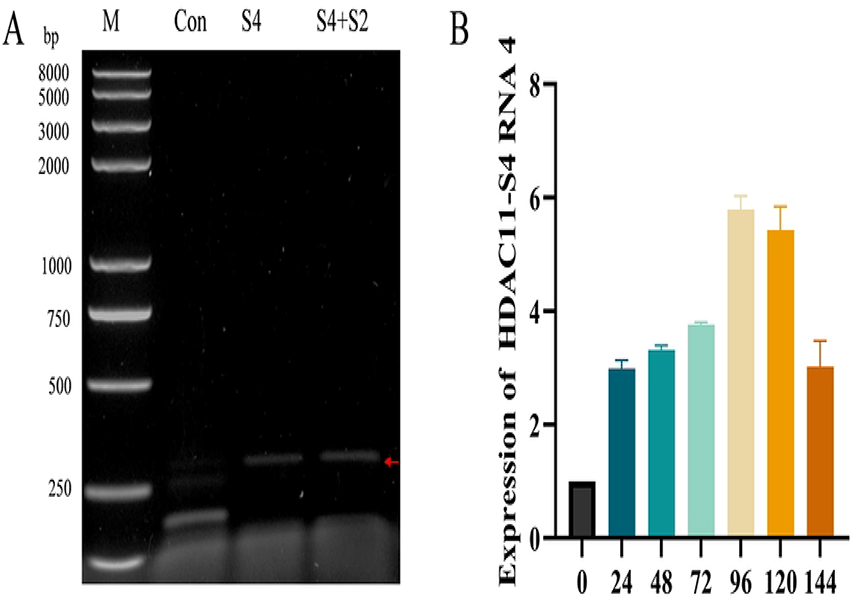
HDAC11-S4 RNA 4 was validated by Northern blotting and in situ hybridization. **A**, Validation of HDAC11-S4 RNA 4 by Northern blotting. ‘18S’ indicates 18S rRNA, the internal control. The total RNA extracted from the BmCPV-infected midgut was separated on 1% agarose-formaldehyde gels and transferred to Hybond-N+ nylon membranes. Northern blotting was conducted with biotin-labeled DNA targeting the junction site of chimeric HDAC11-S4 RNA 4. Total RNA extracted from the non-BmCPV-infected midgut was used as a control. **B**, Validation of HDAC11-S4 RNA 4 by *in situ* hybridization. BmN cells with/without BmCPV infection at 48 h postinfection were digested with proteinase K and hybridized with biotin-labeled DNA targeting the junction site of chimeric HDAC11-S4 RNA 4. Hybridization signals were detected using CY3-labeled streptomycin. Cell nuclei were counterstained with DAPI.

### 1.4 Chimeric HDAC11-S4 RNA 4 is formed in a BmCPV-encoded RdRp-independent manner

The RNA polymerase encoded by sNSVs is required for cap-snatching (Rialdi et al., 2017). To determine whether RdRp encoded by BmCPV is required for the formation of chimeric HDAC11-S4 RNA 4, pIZT-CS4 with the complete cDNA sequence of BmCPV S4 dsRNA (Guo et al., 2018) and a mixture of pIZT-CS4 and pIZT-CS2 with the complete cDNA sequence of BmCPV S2 dsRNA encoding RdRp were transfected into BmN cells, and the total RNAs extracted at 48 h posttransfection were used to evaluate the formation of chimeric HDAC11-S4 RNA 4 by PCR. The desired specific PCR product could be amplified from the two extracted RNA samples, but the specific PCR product could not be obtained from the RNAs extracted from the BmN cells without transfection, suggesting that the formation of chimeric HDAC11-S4 RNA 4 is not mediated by RdRp which is encoded by BmCPV (Figure 5A).

**Figure 5.**
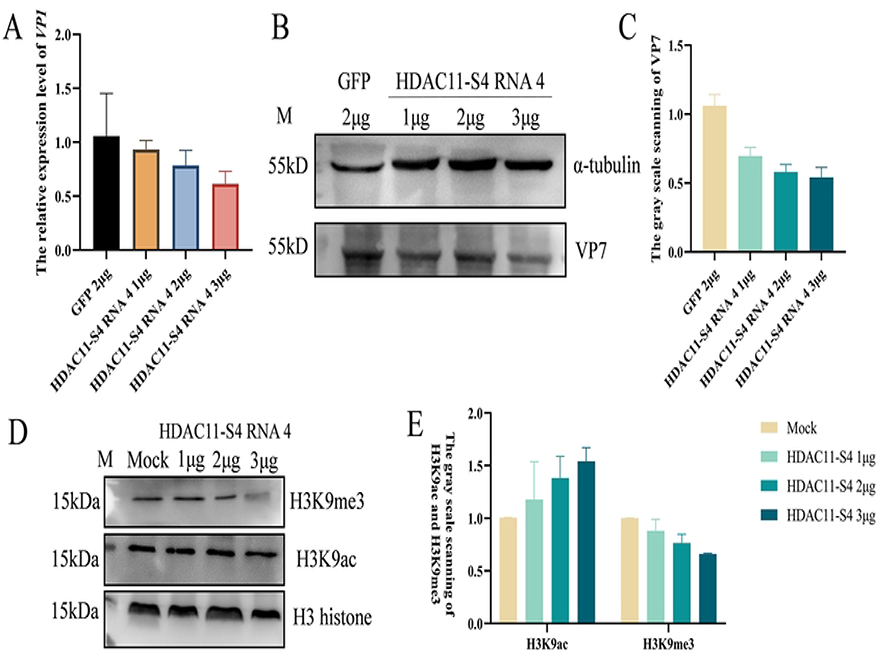
Formation and expression pattern of chimeric HDAC11-S4 RNA 4. **A**, Formation of chimeric HDAC11-S4 RNA 4. pIZT-CS4 and a mixture of pIZT-CS4 and pIZT-CS2 were transfected into BmN cells, and total RNA was extracted at 48 h posttransfection to determine the formation of chimeric HDAC11-S4 RNA 4 with PCR. **B**, Expression pattern of chimeric HDAC11-S4 RNA 4. The total RNAs extracted from the midgut infected with BmCPV at 0-144 h postinfection were used as templates, and the expression level of chimeric HDAC11-S4 RNA 4 was determined by qRT‒PCR. The TIF-4A gene was used as an internal reference.

### 1.5 The level of chimeric HDAC11-S4 RNA 4 increases upon viral infection

To understand the HDAC11-S4 RNA 4 expression pattern, qRT‒PCR was conducted. The results showed that the level of chimeric HDAC11-S4 RNA 4 increased following BmCPV infection, but the expression level decreased at 144 h postinfection (Figure 5B). Moreover, our results showed that chimeric HDAC11-S4 RNA 4 was not detectable in BmCPV virions, indicating that chimeric HDAC11-S4 RNA 4 cannot be packaged into virions.

### 1.6 BmCPV replication is regulated by chimeric HDAC11-S4 RNA 4

To understand the function of chimeric HDAC11-S4 RNA 4, the effect of chimeric HDAC11-S4 RNA on BmCPV replication was investigated. The qRT‒PCR results showed that the *vp*1 gene expression level decreased in chimeric HDAC11-S4 RNA 4-transfected BmN cells in a transfection dose-dependent manner (Figure 6A). Western blot results indicated that VP7 expression was also decreased in chimeric HDAC11-S4 RNA 4-transfected BmN cells compared to green fluorescent protein (GFP) RNA-transfected BmN cells (Figure 6B, 6C), suggesting that BmCPV replication was inhibited.

**Figure 6.**
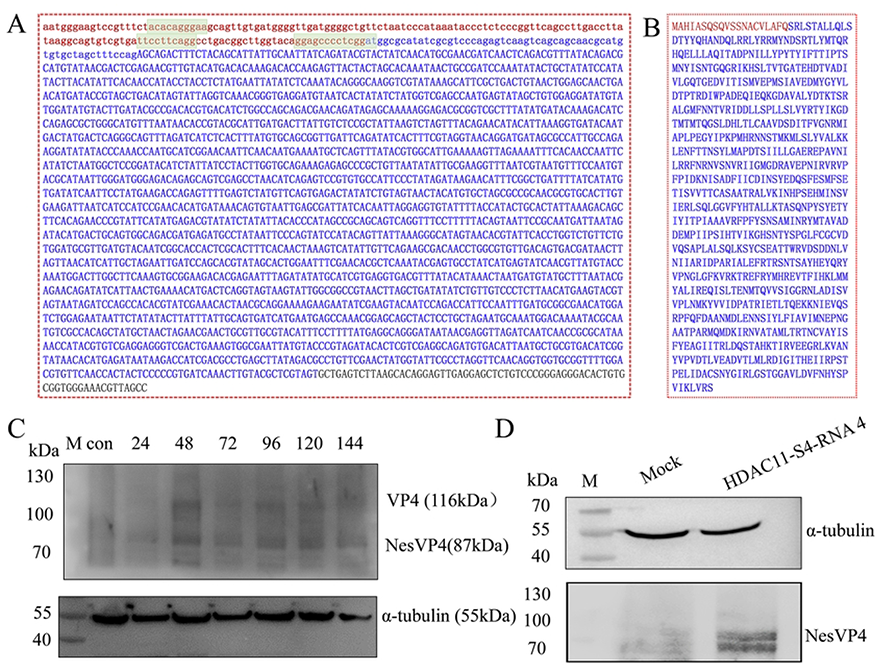
Effect of chimeric HDAC11-S4 RNA 4 on BmCPV viral gene expression and H3K9me3/H3K9ac of histones. **A**, Effect of chimeric HDAC11-S4 RNA 4 on BmCPV *vp1* gene expression levels. BmN cells (10^6^) transfected with 1, 2 or 3 μg of chimeric HDAC11-S4 RNA 4 (HCPV-20-4) were inoculated with BmCPV at 24 h posttransfection, and the relative expression level of the BmCPV *vp1* gene was determined at 48 h posttransfection by qRT‒PCR. gfp RNA (produced by *in vitro* transcription)-transfected cells infected with BmCPV were used as a control. **B**, Effect of chimeric HDAC11-S4 RNA 4 on BmCPV VP7 levels. BmN cells (10^6^) transfected with 1, 2 or 3 μg of chimeric HDAC11-S4 RNA 4 were inoculated with BmCPV at 24 h posttransfection, and the VP7 protein level was determined at 48 h postinfection by Western blotting. gfp RNA (produced by *in vitro* transcription)-transfected cells infected with BmCPV were used as a control. **C**, The grayscale intensity of the Western blot signal bands in Figure 6-B was analyzed by ImageJ software. **D**, Effect of chimeric HDAC11-S4 RNA 4 on H3K9me3 and H3K9ac of histone 3. BmN cells (10^6^) transfected with 1, 2 or 3 μg of chimeric HDAC11-S4 RNA 4 were inoculated with BmCPV at 24 h after transfection, and H3K9me3 and H3K9ac of histone 3 were determined at 48 h postinfection by Western blotting with anti-3K9me3 and anti-H3K9ac antibodies. Histone 3 was used as an internal reference. Mock lane, cells transfected with the *gfp* gene. 1 μg, 2 μg and 3 μg lanes, the cells transfected with 1 μg, 2 μg or 3 μg of HDAC11-S4 RNA 4. **E**, The grayscale intensity of the Western blot signal bands in Figure 6-D was analyzed by ImageJ software.

### 1.7 H3K9me3/H3K9ac of histone 3 is regulated by chimeric HDAC11-S4 RNA 4

Chimeric HDAC11-S4 RNA 4 was a hybrid molecule consisting of the HDAC11 gene transcript and a fragment of the sense RNA of BmCPV S4 dsRNA. Therefore, the effect of chimeric HDAC11-S4 RNA on the methylation and acetylation of histone H3 was determined by Western blotting. The results showed that the trimethylation of histone 3 lysine 9 (H3K9me3) was decreased and the acetylation of histone 3 lysine 9 (H3K9ac) was increased in chimeric HDAC11-S4 RNA-transfected BmN cells compared with BmN cells transfected with gfp RNA (Figure 6D, E).

### 1.8 Chimeric HDAC11-S4 RNA 4 encodes a truncated viral VP4 with N-terminal extensions derived from host HDAC11 RNA

To explore whether chimeric HDAC11-S4 RNA has the potential to encode proteins, ORF finder (https://www.ncbi.nlm.nih.gov/orffinder/) was used to predict the ORF of chimeric HDAC11-S4 RNA-4. The results showed that the chimeric RNA may encode a novel protein (termed NesVP4) consisting of is a truncated viral VP4 (768 amino acid residues) with N-terminal extensions (20 amino acid residues) derived from host HDAC11 RNA (Figure 7A, 7B). To confirm this result, the BmCPV-infected midgut was analyzed in different infection phases by Western blotting with an anti-VP4 antibody, and two specific signal bands representing VP4 (116 kDa) and NesVP4 (87 kDa) were observed, implying that NesVP4 was translated by the chimeric RNA (Figure 7C). Moreover, the specific signal band representing NesVP4 could be observed in the cells transfected with HDAC11-S4 RNA 4 obtained by *in vitro* transcription (Figure 7D). These results indicated that NesVP4 was translated by HDAC11-S4 RNA 4.

**Figure 7.**
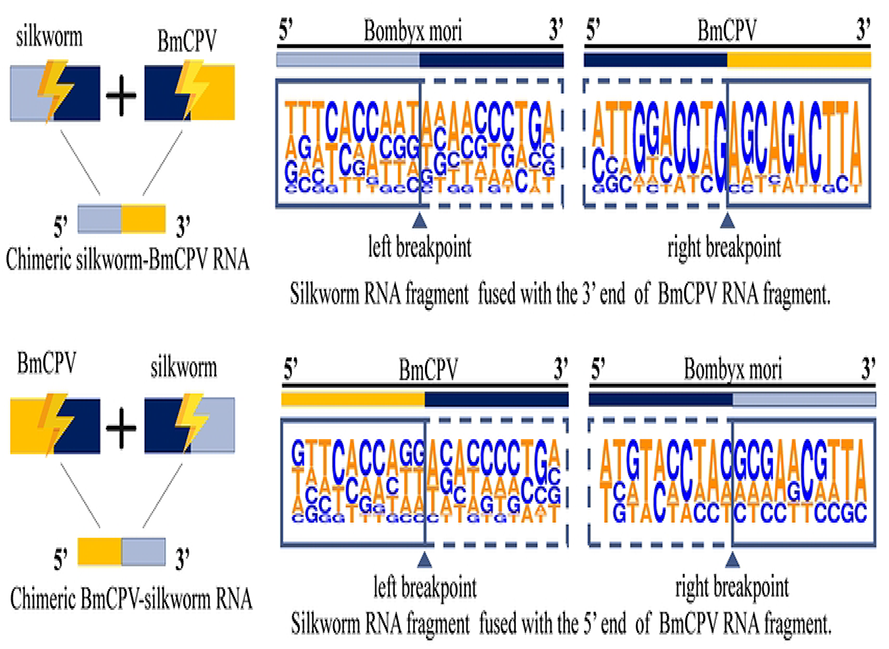
Chimeric HDAC11-S4 RNA 4 encodes a truncated viral VP4 with N-terminal extensions derived from host HDAC11 RNA. **A**, Chimeric HDAC11-S4 RNA 4 sequence. Lowercase letters represent the HDAC11 gene mRNA fragment with HDAC11-S4 RNA 4. Uppercase letters represent the BmCPV S4 RNA fragment with HDAC11-S4 RNA 4. Blue letters represent the speculated open reading frame, and the box represents the hypothesized IRES-like sequence. **B**, Deduced sequence of the NesVP4 protein encoded by chimeric HDAC11-S4 RNA 4. Red letters represent the amino acid sequence derived from HDAC11 mRNA, and blue letters represent the amino acid sequence derived from sense RNA of the BmCPV S4 segment. **C**, Identification of NesVP4 encoded by chimeric HDAC11-S4 RNA in the BmCPV-infected midgut by Western blotting. Proteins from the midguts of silkworms infected with BmCPV at 24-144 h postinfection were separated by SDS‒PAGE. After the proteins on the gel were transferred to a nitrocellulose membrane, Western blotting was performed. An anti-VP4 antibody was used as a primary antibody, and HRP-conjugated goat anti-mouse IgG was used as a secondary antibody. α-Tubulin was used as an internal reference. Lane M, molecular marker; con lane, midgut not infected with BmCPV; lane 24-lan144, midgut infected with BmCPV at 24-144 h postinfection with BmCPV. **D**, Identification of NesVP4 in BmN cells transfected with chimeric HDAC11-S4 RNA 4 by Western blotting. Lane M, molecular marker; Mock lane, cells without transfection; HDAC11-S4 RNA 4 lane, BmN cells transfected with HDAC11-S4 RNA 4 at 48 h.

## 2 Discussion

Previous studies have indicated that chimeric RNAs can be generated during the infection of sNSVs, HIV (single-stranded RNA virus), and SV40/JCV/HSV-1/ADV (double-stranded DNA viruses) (Clohisey et al., 2020; Eul and Patzel, 2013; Ho et al., 2020; Lee et al., 2022; Toba et al., 2022). In this study, chimeric silkworm-BmCPV RNAs were identified in the BmCPV-infected midgut. BmCPV is a segmented dsRNA virus belonging to the *Cypovirus* genus of the Reoviridae family (Cui et al., 2019). To our knowledge, this is the first report showing that chimeric silkworm-BmCPV RNAs can be formed during BmCPV infection.

Previous studies have shown that in cells infected with DNA viruses that integrate into the host genome (e.g., hepatitis B virus or human papillomavirus), chimeric RNAs that are partly mapped to host RNA and partly mapped to viral RNA can be formed because the host genomic DNA region integrating viral DNA is transcribed (Brant et al., 2018; Sung et al,, 2012; Zhao et al., 2016). BmCPV is a dsRNA virus that replicates in the cytoplasm and does not have a nuclear phase in its life cycle. To date, no event in which viral cDNA is integrated into the silkworm genome has been identified. Therefore, the chimeric silkworm-BmCPV RNAs found in this study were not produced by the integration of cDNAs originating from BmCPV RNAs into the silkworm genome and subsequent transcription.

In the process of sNSV infection, chimeric host‒virus RNAs with highly diverse host-derived 5’ sequences can be generated by a cap-snatching mechanism, and the RdRp encoded by sNSVs is required for the formation of chimeric RNAs (Ho et al., 2020; Olschewski et al., 2020; Kuang et al., 2022). sNSVs perform cap-snatching in the nucleus, while bunyaviruses perform cap-snatching in the cytoplasm. The bunyavirus L protein is suggested to be responsible for cap binding and cleavage of host mRNA to generate a capped RNA fragment. Subsequently, the capped RNA fragment is used as a primer for viral transcription to form a chimeric mRNA (Olschewski et al., 2020). In this study, we found that the fragments derived from silkworm RNA could fuse with the 5’ or 3’ terminus of the viral RNA fragment and that the RdRp encoded by BmCPV was not required for the formation of HDAC11-S4 RNA 4. Therefore, we hypothesized that chimeric silkworm-BmCPV RNAs were formed via a mechanism different from the cap-snatching mechanism.

It has been reported that some chimeric mRNAs can be generated by readthrough transcription and aberrant splicing during HSV-1 (Depledge et al. 2019) and HIV (Lee et al., 2022) infection, respectively. The BmCPV genome consists of 10 segmented dsRNAs, and each segmented dsRNA forms a transcript, suggesting that the chimeric virus‒virus RNAs found in this study were not formed by readthrough transcription.

Cis-splicing is a splicing reaction that removes introns and joins the exons included within the same RNA transcript in the nucleus to form mature RNA. Trans-splicing is a splicing reaction between two RNA molecules. Previous studies have shown that chimeric RNAs, including hybrid molecules between viral RNA and host RNA or viral RNA and viral RNA, can be generated by trans-splicing during infection by ADV, HIV, SV40 and JCV (Kikumori et al., 2002; Caudevilla et al., 2001; Eul et al., 2013; Saribas et al., 2018). Usually, the sequences of the splice sites carried by RNA molecules must be located close to the consensus splice site sequences (for 5′ splice site: GU; for 3′ splice site: AG), and partial sequences in the introns of one RNA molecule complement intron regions of another RNA molecule (Lasda and Blumenthal, 2011; Hou et al., 2022). CPV replication and gene transcription occur in the cytoplasm, and consensus splice site sequences matching the trans-splicing mechanism for the formation of chimeric silkworm-BmCPV RNAs were not found in the corresponding viral RNAs and silkworm RNAs. Whether chimeric BmCPV/silkworm-BmCPV RNAs are formed via a trans-splicing mechanism is worthy of further study. Moreover, it has been indicated that nonhomologous recombination and homologous recombination can occur between different RNA molecules (Chetverin et al., 1999). A high frequency of homologous recombination is observed by some RNA viruses in a replicative template switch mechanism. RNA‒RNA recombination has been found between viral RNAs and even between viral RNAs and host RNAs (Sztuba-Solińska, et al., 2011; Zou et al., 2019; Wang et al., 2022). Therefore, we cannot exclude the possibility that chimeric BmCPV/silkworm-BmCPV RNAs are generated by RNA recombination.

It has been reported that host‒virus chimeric RNAs identified by the RNA sequencing of cells infected with severe acute respiratory syndrome coronavirus 2 are likely artifacts arising from random template switching of reverse transcriptase and/or sequence alignment errors (Yan et al., 2021). To eliminate misjudgment caused by sequencing artifacts, reverse transcriptional noise, template-switching and scrambled junctions of RNAs, chimeric HDAC11-S4 RNA 4 was selected for further validation. The authenticity of chimeric HDAC11-S4 RNA 4 was further confirmed by Northern blotting and cell *in situ* hybridization.

It has been indicated that some chimeric host‒virus mRNAs can be translated into chimeric proteins, viral proteins with host-encoded extensions or new host‒virus-encoded proteins using uAUGs within cap-snatched host transcripts as start codons (Ho et al., 2020). In this study, two specific signal bands, one representing viral VP4 (116 kDa) and the other representing a novel protein (termed NesVP4) consisting of a truncated viral VP4 with N-terminal extensions (20 amino acid residues) derived from host HDAC11 RNA, suggesting that chimeric HDAC11-S4 RNA 4 can be translated into a protein using an AUG codon within the host RNA sequence of chimeric HDAC11-S4 RNA as the start codon. BmCPV VP4, which shows RNA guanylyltransferase activity, is a viral turret protein encoded by the BmCPV S4 segment (Cao et al., 2012), and 75 amino acid residues at the N-terminus of VP4 are required for embedding viral particles into polyhedrons (Ikeda et al., 2006). During virus assembly, the possibility that VP4 could be replaced by NesVP4 encoded by chimeric HDAC11-S4 RNA to form a defective virus is worthy of further study.

It has been reported that the host translational machinery is recruited to chimeric host‒virus mRNAs generated via a cap-snatching mechanism, which results in global host transcriptional suppression and evasion of antiviral surveillance by RIG-I-like receptors (Decroly et al., 2011). Novel proteins translated by chimeric host‒virus mRNAs impact viral replication and viral virulence and drive host T-cell immunity (Russell, 2020; Wang et al., 2020). In this study, we found that viral gene expression decreased, H3K9me3 marks decreased and H3K9ac marks increased in cells transfected with chimeric HDAC-VP4 RNA 4, suggesting that viral replication and host gene expression can be regulated by chimeric HDAC-VP4 RNA. In short, our results indicated that during BmCPV infection, a novel mechanism different from that described in previous reports allows the genesis of chimeric silkworm-BmCPV RNAs with biological functions.

## 3 Materials and methods

### 3.1 RNA_seq

Newly molted 5th-instar silkworm larvae (Jingsong Strain) were fed mulberry leaves coated with 10^8^ BmCPV polyhedra/ml for 8 h, followed by feeding with fresh leaves at 25°C. The midguts of silkworms infected with BmCPV were dissected at 48, 96 and 144 h postinoculation.

Total RNA was isolated from collected midguts with RNeasy R Plus Mini Kits (Qiagen, Valencia, CA, USA). After genomic DNA was digested by RNase-free DNase (Qiagen, Valencia, CA, USA), the quantity of RNA was determined with a NanoDrop 2000 Spectrophotometer (Thermo Scientific, Wilmington, USA). RNA integrity was evaluated with an Agilent 2100 Bioanalyzer (Agilent Technologies, Palo Alto, CA, USA), and RNA integrity numbers (RINs) were determined based on agarose gel electrophoresis. After ribosomal RNA was removed with an Epicenter Ribo-zero™ kit (Epicenter, Charlotte, NC, USA), the RNA was fragmented into 200-300 nt fragments with fragmentation buffer (New England Biol, Peking, China), followed by reverse transcription with random hexamers to generate the first strand of cDNA. Finally, cDNA libraries were constructed using PCR.

The library was quantified with a Qubit 2.0 Fluorometer and diluted to 1.5 ng/μL. The quality of the cDNA library was assessed in an Agilent 2100 Bioanalyzer. Furthermore, the effective concentration was determined by quantitative PCR. The complete library was sequenced by Novogen. (Peking, China) on a HiSeq 2500 Sequencer (Illumina, San Diego, CA, USA). All sequencing data were deposited in the NCBI database under accession numbers SRR22891215, SRR22891214 and SRR22891213.

### Preprocessing of RNA_Seq data

Illumina Casava 1.8 base-calling software was used to convert the image data obtained by the high-throughput sequencer into sequence data. The FastQC toolkit (http://www.bioinformatics.babraham.ac.uk/projects/fastqc/) was used to evaluate the quality of raw reads according to the QPhred score. After removing sequences related to adaptors with SeqPrep software (https://github.com/jstjohn/SeqPrep), low-quality reads (QPhred score < 20) were trimmed, and reads that were shorter than 50 bp and or contained N bases (unknown bases) were removed using Sickle software (https://github.com/najoshi/sickle). rRNA reads were removed by aligning reads to the SILVA SSU (16S/18S) and SILVA LSU (23S/28S) databases using SortMeRNA software (http://bioinfo.lifl.fr/RNA/sortmerna/) to generate high-quality reads.

### 3.3 De novo assembly and prediction of chimeric silkworm-viral RNA

StringTie software (1.2.0) (Pertea et al., 2015) was applied to assemble the obtained effective reads to generate transcripts using the *B. mori* genome (https://silkdb.bioinfotoolkits.net/_resource/Bombyx_mori/download/chromosome.fa.tar.gz.) and BmCPV genomic dsRNAs (GU323605, GQ924586, GQ924587, GU323606, GQ294468, GQ294469, GQ150538, GQ150539, GQ924588, and GQ924589) as references. Subsequently, STAR-Fusion was used to analyze fusion transcripts (Haas et al., 2019), and the obtained fusion transcripts were visualized by generating a Circos diagram with R software.

### 3.4 RT‒PCR and Sanger sequencing

Total RNA was isolated from collected BmCPV-infected midguts with RNeasy R Plus Mini Kits (Qiagen, Valencia, CA, USA). After genomic DNA was digested by RNase-free DNase (Qiagen, Valencia, CA, USA), the RNA was reverse transcribed into cDNA. Furthermore, PCR was conducted with primers (Table S2) designed according to the flanking sequences of the junction sites of predicted chimeric silkworm-viral RNAs. The PCR products were cloned into pMD-18T for sequencing.

### 3.5 Northern blotting

To further confirm the authenticity of chimeric HDAC11-S4 RNA 4, the total RNAs extracted from midguts infected with BmCPV were identified by Northern blotting using a commercial kit (Ambion, Austin, TX, USA). Briefly, total RNA (50 μg) was separated on 1% agarose–formaldehyde gels and transferred to Hybond-N+ nylon membranes (Roche, Basel, Switzerland). Northern blotting was conducted with biotin-labeled DNA (bio-CTGTAGAAAGTCTGCTGATCGATACCGCGACG, synthesized by Sangon Biotech, Shanghai, China) targeting the junction site of chimeric HDAC11-S4 RNA 4. Signals were visualized with a Biotin Chromogenic Detection Kit (Thermo Scientific, Waltham, MA, USA).

### 3.6 *In situ* hybridization

A total of 1×10^4^ cultured BmN cells infected with BmCPV in 24-well plates at 48 h postinfection were digested with proteinase K and fixed in 4% formaldehyde at 4°C for 1 h. The cells were hybridized with the biotin-labeled DNA probe mentioned above at 37°C overnight according to the manual of the Ribo™ Fluorescent *in situ* Hybridization Kit (China, BOSTER, Cat: MK1030). CY3-labeled streptomycin was used to visualize hybridization signals. Nuclei were stained with 4,6-diamidino-2-phenylindole (DAPI). Cell images were captured using a Leica DM2000 microscope (Leica, Wetzlar, Germany).

### 3.7 Cell transfection

A total of 1×10^6^ cultured BmN cells in 6-well plates were transfected with pIZT-CS4 (2 μg) containing the complete cDNA sequence of BmCPV S4 dsRNA (Guo et al., 2018) or a mixture of pIZT-CS4 (1 μg) and pIZT-CS2 (1 μg) containing the complete cDNA sequence of BmCPV S2 dsRNA encoding RdRp using Roche-X Gem (Switzerland, Roche, Cat: 6366236001). At 48 h posttransfection, the cells were collected for the extraction of total RNA.

### 3.8 Characterization of the expression pattern of chimeric HDAC11-S4 RNA 4

To characterize the expression pattern of chimeric HDAC11-S4 RNA 4, the total RNAs extracted from midguts infected with BmCPV at 0, 24, 48, 72, 96, 120 and 144 h postinoculation were reverse transcribed into cDNA using random primers (First Strand cDNA Synthesis Kit, Transgene, Beijing, China). The expression levels of chimeric HDAC11-S4 RNA 4 were determined by real-time PCR using a pair of specific primers, qHCPV20-4 (Table S2). The translation initiation factor eIF-4A (TIF-4A) gene was utilized as an internal reference.

### 3.9 Plasmid construction and transcription in vitro

The cDNA sequence of chimeric HDAC11-S4 RNA 4 (Figure 7A) controlled by the T7 promoter was chemically synthesized and cloned into the *Eco*RI and *Hin*dIII sites of the pUC57 vector to generate pUCHCPV-20-4. After being digested by HindIII, linearized pUCHCPV-20-4 was used as a template for *in vitro* transcription by T7 RNA polymerase (TaKaRa, Dalian, China) to generate chimeric HDAC11-S4 RNA 4. After removing the plasmid DNA with DNaseI, chimeric HDAC11-S4 RNA 4 was purified by extraction with phenol/chloroform treatment and precipitation with ethanol.A similar method was used to generate *gfp* RNA.

### 3.10 Effect of chimeric HDAC11-S4 RNA 4 on BmCPV viral gene expression and histone modification

First, 1×10^6^ BmN cells (1 ml) were transfected with chimeric HDAC11-S4 RNA 4 (1, 2 or 3 μg), and 24 h later, the cells were inoculated with BmCPV (MOI=2). The collected cells at 48 h postinoculation were used for detection of viral gene expression and modification of histone 3. For viral gene expression, the transcription level of the *vp1* gene was determined by qRT‒PCR with CPV-S1 primers (Table S2). The levels of VP7, H3K9me3 and H3K9ac were determined by Western blotting with corresponding antibodies. Histone 3 was used as an internal reference. Cells transfected with *gfp* RNA were used as a control.

## Acknowledgments

This research was supported by the National Natural Science Foundation of China (32072792, 31872424, and 31972620) and the Priority Academic Program of Development of Jiangsu Higher Education Institutions. The funders had no role in the study design, data collection, analysis, decision to publish, or manuscript preparation.

---

**Figure S1.**
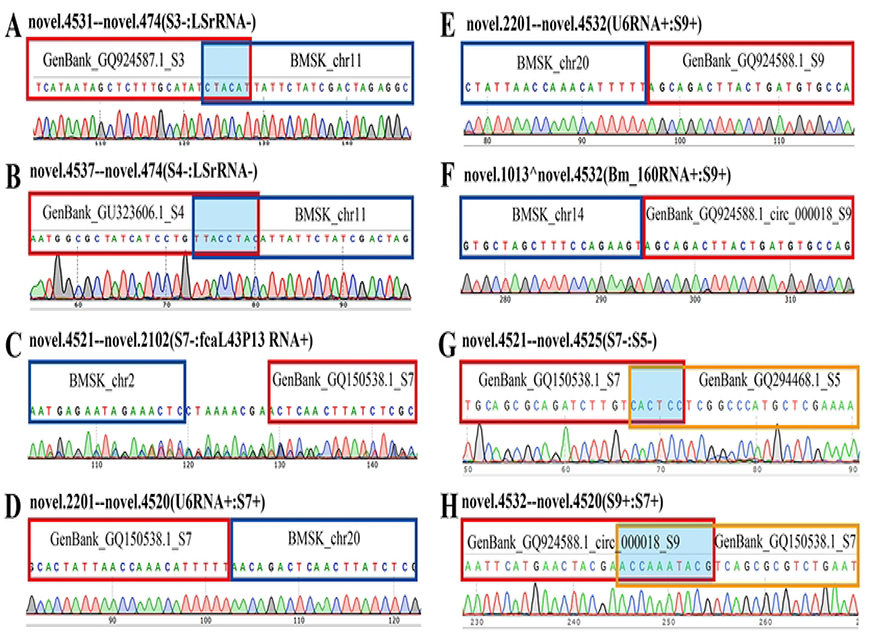
The flanking sequences of the breakpoints of left/right parental RNAs.

**Table S1 Information for chimeric silkworm-BmCPV RNAs identified in the midguts of silkworms infected with BmCPV.**

**Table S2 The primers used in this study.**

